# Trehalose as an alternative to glycerol as a glassing agent for in vivo DNP MRI

**DOI:** 10.1101/866665

**Authors:** Jeffrey R. Brender, Shun Kishimoto, Gareth R. Eaton, Sandra S. Eaton, Yu Saida, Murali C. Krishna

## Abstract

In dynamic nuclear polarization (DNP), the solutions of the hyperpolarizable molecule and the paramagnetic agent need to form a glass when frozen to attain significant levels of polarization in reasonable time periods. Molecules which do not form glasses by themselves are often mixed with excipients to form glasses. While glassing agents are often essential in DNP studies, they have the potential to perturb the metabolic measurements that are being studied. Glycerol, the glassing agent of choice for in vivo DNP studies, is effective at reducing ice crystal formation during freezing but is rapidly metabolized, potentially altering the redox and ATP balance of the system. As a biologically inert alternative to glycerol, we show here that 15–20 wt % trehalose yields a glass that polarizes samples more rapidly than the commonly used 60% wt formulation of glycerol and yields similar polarization levels within clinically relevant timeframes. Trehalose may be an attractive alternative to glycerol for situations where there may be concerns about glycerol’s glucogenic potential and possible alteration of the ATP/ADP and redox balance.

## Introduction

Dissolution dynamic nuclear polarization DNP has opened an unique window into cellular metabolism by allowing the non-invasive tracking of exogenous metabolic tracers.^*1*, *2*^ By probing the activity of specific targeted enzymes rather than measuring absolute concentrations of metabolites, dissolution DNP has an advantage in isolating specific metabolic activities over static techniques such as MRSI, which measure an equilibrium state that may stem from many processes. However, the widespread use of dissolution DNP is limited by the availability of suitable metabolic tracers. Most dissolution DNP studies to date have used pyruvic acid as a tracer. Pyruvate’s dominance in DNP research is partly due to the central position pyruvate occupies in glucose metabolism,^*3*^ but also due to its physical properties. In contrast to most other metabolites, pyruvate is a liquid at room temperature. The liquid state of pyruvic acid ensures the radical is evenly mixed throughout the sample. In a solid sample dissolved in water, phase separation can occur during freezing.^*4*, *5*^ This is problematic as transfer efficiency from the radical to the nuclei of interest in dynamic nuclear polarization is steeply non-linear with respect to the radical- nuclear spin distance.^*6*, *7*^ Clustering of the radical species due to phase separation increases the median radical- nuclear spin distance and may result in sharply reduced transfer efficiencies and poor signal in DNP.

To reduce phase separation during freezing, glassing agents are commonly employed in studies of non-self-glassing tracers.^*8*^ Glycerol has emerged as the glassing agent of choice for *in vivo* studies,^*5*^ due to its favorable safety profile^*9*^ and relatively high glassing efficiency. Although glycerol is safe at the concentrations used in DNP studies,^*9*^ it is not metabolically inert.^*10*^ Glycerol is rapidly transported into the cell by aquaporin^*11*^ and is readily metabolized to glyceraldehyde-3-phosphate, consuming ATP and FAD in the process. It may therefore potentially perturb measurements of the TCA cycle by the consumption of ATP during the phosphorylation to glycerol-3-phosphate and by alteration of the FAD/FADH_2_ and NAD/NADH cytosolic redox balance from the glycerol-3-phosphate shuttle.

An improved glassing agent with a favorable safety profile that does not affect metabolism would help expand the clinical translation of DNP beyond pyruvate to other metabolic tracers. Trehalose is a non-reducing disaccharide commonly used as a pharmaceutical excipient to stabilize proteins in an immobile glass matrix.^*12*^ Trehalose has an unusually high glass transition temperature on account of its anisotropic hydrogen bonding pattern,^*13*^ which inhibits ice nucleation^*13*^ and favors the formation of amorphous glasses over ordered crystalline solids or liquids. Clinical tests of trehalose IV injections have not shown side effects in concentrations as high as 9% wt%, far exceeding the 0.5-1% range expected in dissolution DNP after the filtration step. In contrast to glycerol, trehalose is only metabolized in the small intestine where trehalase is present and is therefore metabolically inert in IV injections (less than 0.5% is absorbed by passive diffusion in humans).^*14*^ We show here that glasses containing ~15% trehalose give polarizations similar to the commonly used glycerol formulation, but at much faster polarization rates.

## Methods

### Dynamic Nuclear Polarization

4.5 M [1-^13^C]urea or 3M [1-^13^C]glycine (30◻μL), containing 15◻mM Oxo63, and the indicated amounts of glassing agents, was hyperpolarized at 3.35◻T and 1.4◻K using the Hypersense DNP polarizer (Oxford Instruments, Abingdon, UK) according to the manufacturer’s instructions. Polarization buildup rates were calculated assuming single exponential growth. For the mouse imaging experiment, the hyperpolarized urea was rapidly dissolved in 4.5◻mL of a superheated HEPES based buffer before being intravenously injected through a catheter placed in the tail vein of the mouse (12◻ μL/g body weight). Spectra were recorded on a 3T MRSolutions scanner.

### EPR Studies

Electron spin relaxation times of 15 mM OXo63 trityl samples in water:trehalose solutions were measured at 5K using a Bruker E580 pulsed EPR spectrometer with the sample cooled in a He gas atmosphere using an Oxford CF935 cryostat and a Bruker/ColdEdge “Stinger” closed cycle He cooling system. With trehalose present, T_m_ and T_1_ were measured by spin-echo methods.^*15*^ T1 values were calculated from UPEN analysis^*16*, *17*^ of inversion recovery experiments using 100 exponentials without variable smoothing and an additional non-negativity constraint with most measurements made near the peak of the field-dependent spectrum. The signal detection gate encompassed about half of the width of the echo and FID measurements integrated most of the FID intensity. Without trehalose, the EPR signal exhibited an FID which could not be suppressed with attempts to decrease the magnetic field homogeneity, so an echo could not be observed for this sample, and T_m_ was not measured. The absorption spectrum was obtained by field-swept FID detection, followed by calculation of the power spectrum. Samples of OXo63 (sodium salt) and trehalose in water were prepared gravimetrically with the OXo63 concentration kept at 15 mM. Samples containing trehalose were frozen by cooling the 4 mm o.d. quartz tube in liquid nitrogen before inserting it into the cooled cryostat. The sample without trehalose was placed in a 1.1 mm i.d. Teflon tube inside the 4 mm o.d. quartz tube to prevent breaking the tube and inserted into the cold cryostat in an He atmosphere without prefreezing. Air was not removed from the samples before freezing, as previous studies found no effect of air on spin lattice relaxation of trityls in frozen solution.

## Results

To test the potential of trehalose as a biocompatible glassing agent, we recorded the polarization buildup curves of 4.7 M ^13^C urea, a potential marker for perfusion in patients with compromised renal function who cannot tolerate gadolinium or iodinated contrast agents,^*18*^ in the presence of increasing amounts of trehalose. In the absence of trehalose, polarization of urea was slow and the buildup of polarization was negligible in clinically relevant timeframes (30 minutes or less) (Figure 1A orange line). Imaging of a mouse leg xenograft with hyperpolarized urea without trehalose yielded only noise with no discernable signal whatsoever (Figure 2A).

**Figure 1.**
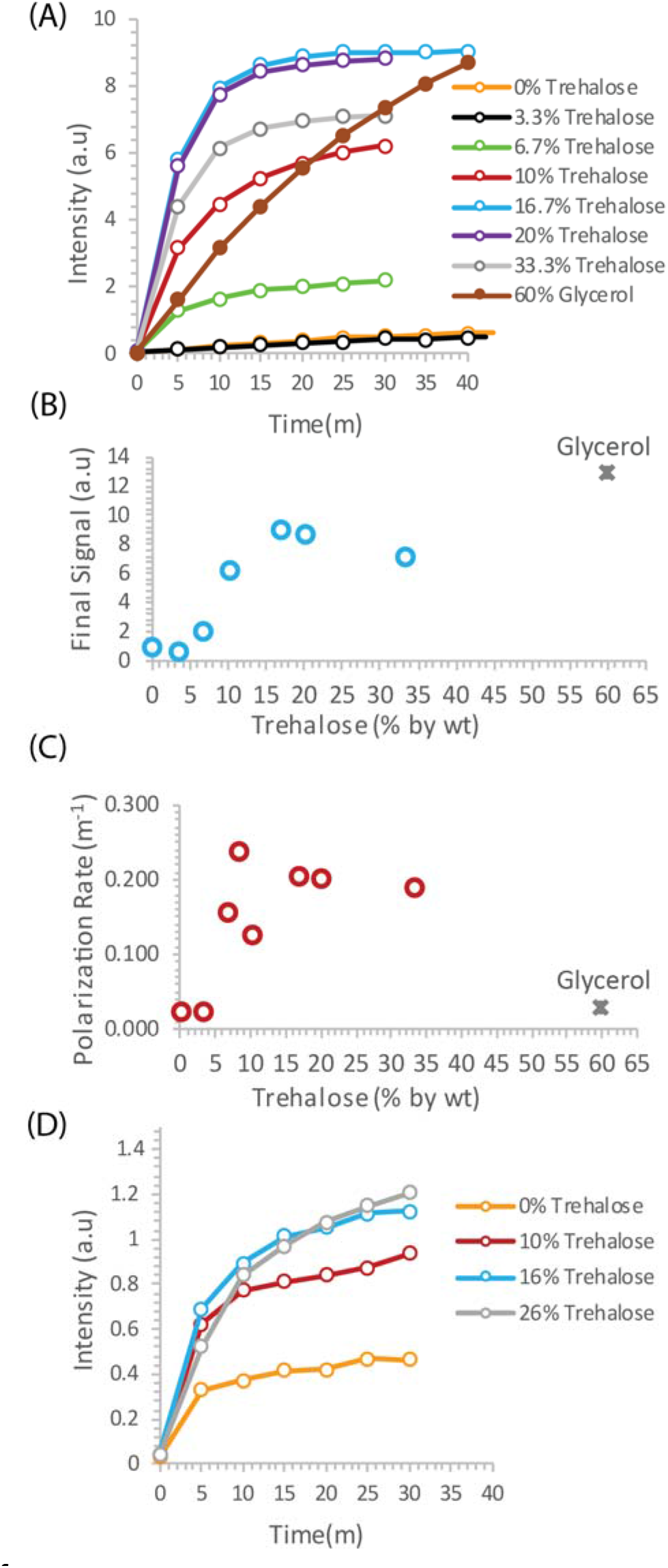
**(A)** DNP buildup of 4.7 M ^13^C urea in trehalose solutions vs. the current 60% glycerol protocol **(B** and **C)** Plot of the final signal **(B)** and polarization rate **(C)** as a function of the trehalose concentration. **(D)** DNP buildup of 3 M ^13^C glycine with increasing concentrations of trehalose as a glassing agent

**Figure 2.**
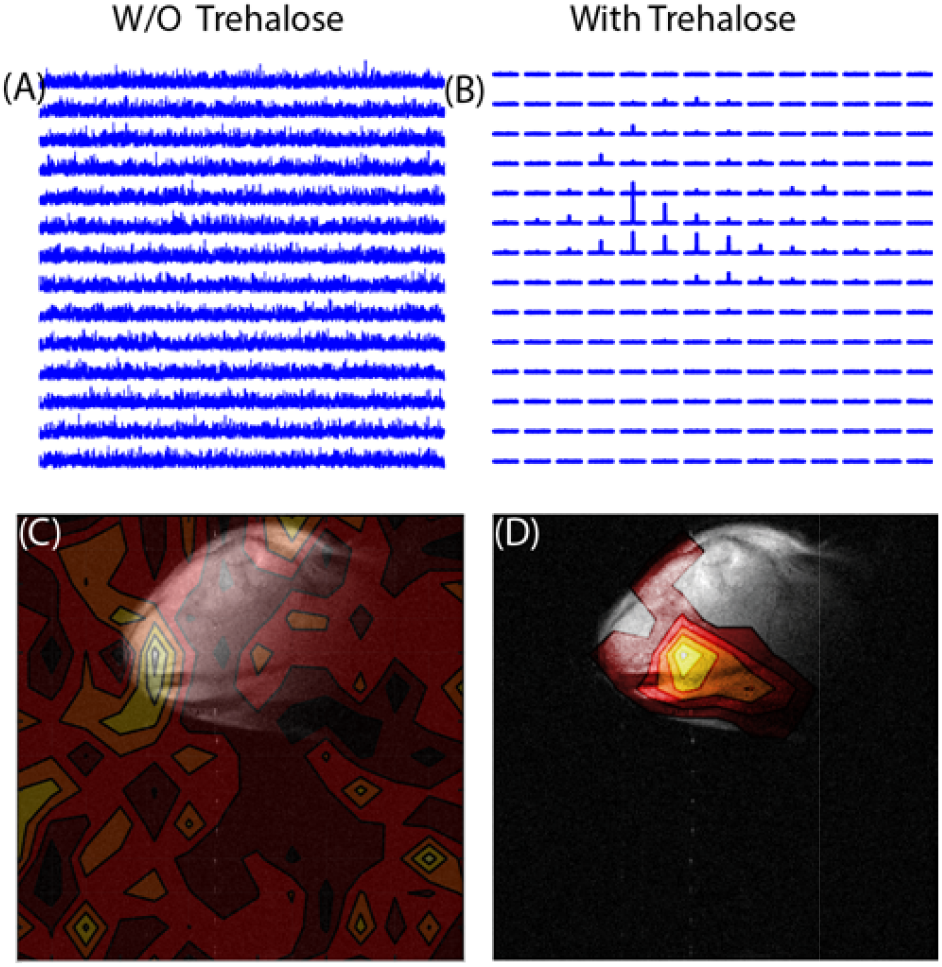
**(A and C)** Imaging of a mouse leg xenograft by CSI imaging with hyperpolarized ^13^C urea in the absence of a glassing agent **(B and D)** Th same mouse leg imaged using 15% trehalose as a glassing agent. A clear signal is evident within the tumor only when trehalose is used as a glassing agent.

The addition of trehalose as a glassing agent markedly increased both the equilibrium polarization and buildup kinetics. Starting at 6.7%, the final signal increases rapidly with concentration before leveling off at approximately 15% trehalose. A similar concentration dependence is also seen with the buildup rate (Figure 1C). In comparison to the standard glycerol preparation, trehalose solutions build up polarization much more quickly but have a lower equilibrium polarization. The increase in polarization led to greatly enhanced imaging *in vivo*. While the urea image without a glassing agent was completely noise, the addition of 20% trehalose in the polarization mixture results in a clear image with urea localized within the tumor (Figure 2B). To confirm that the results are not confined to urea, we repeated the polarization experiments with another difficult to polarize, non-self glassing substrate, ^13^C labelled glycine, with similar results (Figure 1D).

Although trehalose is often cited as having unique molecular properties^*19*–*23*^ that give it anomalous glass forming and ice breaking^*24*–*26*^ capabilities, the molecular properties of trehalose have been the subject of considerable contention,^*27*, *28*^ and other studies have suggested trehalose is not unique among carbohydrates in this regard.^*29*^ To evaluate other potential carbohydrate glassing agents besides trehalose, we measured the polarization buildup of urea in the presence of equivalent amounts of analogous mono- and disaccharides (Figure 3). Trehalose was not unique in this context; fructose, glucose, and PEG400 all had slightly higher equilibrium polarization at an equivalent wt %. Sucrose had significantly lower equilibrium polarization but similar build up kinetics. In comparison, 20% trehalose was an effective compromise between fast buildup and equilibrium polarization, allowing efficient polarization within a clinically relevant timeframe.

**Figure 3.**
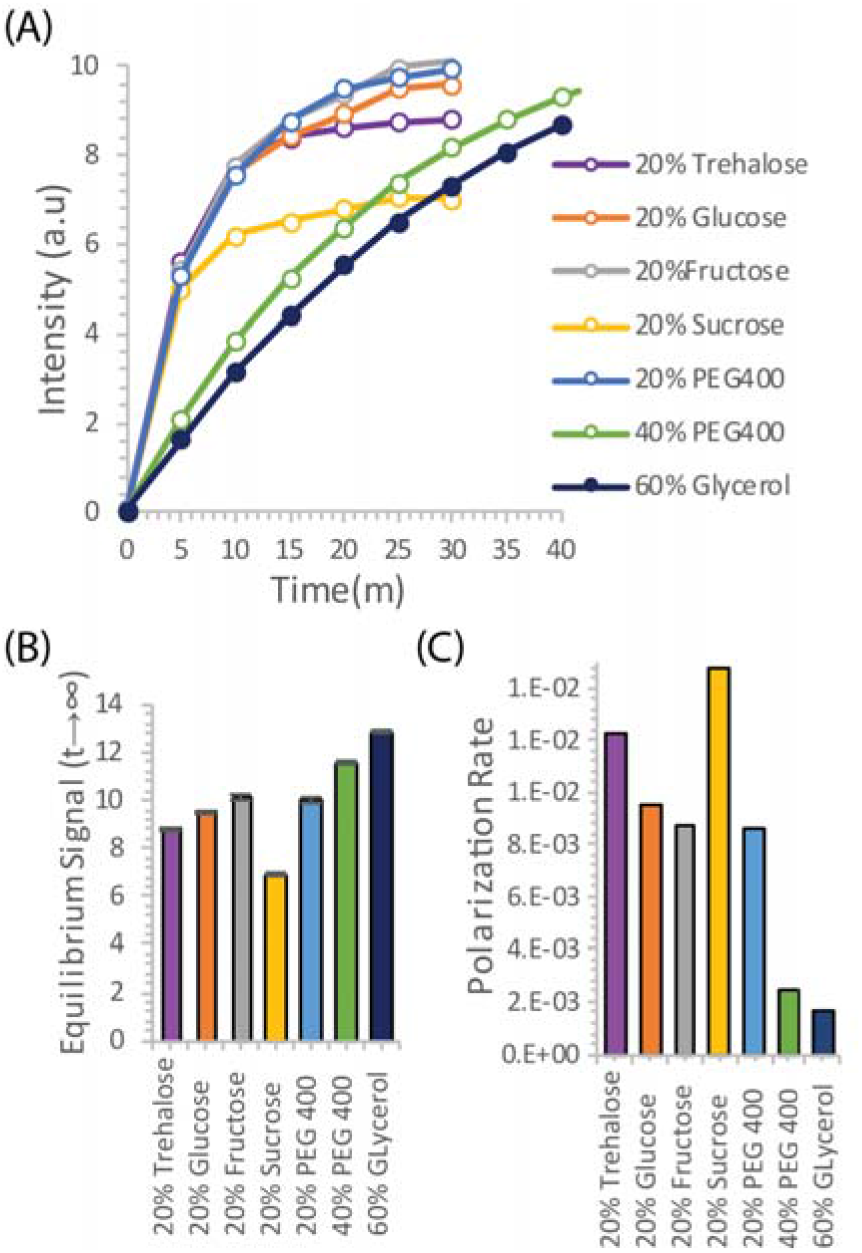
**(A)** DNP buildup of 4.7 M ^13^C urea using trehalose solutions as a glassing agent vs equivalent polyalcohols. **(B** and **C)** Plot of the final signal **(B)** and polarization rate **(C)** for the different polyalcohols

The complex dependence of the polarization efficiency on the trehalose concentration in Figure 1 suggests a possible change in the microenvironment of the radical through a phase separation during freezing or other process. To gain a greater understanding of how trehalose may affect changes in the microenvironment of the Oxo63 radical during the DNP process, we recorded the EPR spectra (figure 4 and spin-lattice (T_1_) and phase memory dephasing (T_m_) relaxation times (Figure 4) of Oxo63 at 5K as a function of trehalose concentration. With roughly ten to sixty times more trehalose than OXo63 and the overall high concentrations of spins, it is likely that there are multiple environments for the radicals, some of which may be rather close neighbors of other radicals. While the complexity of the system makes defining a quantitative physical model difficult, the differences in the microenvironment will likely be reflected in the T_1_ relaxation times, as the most isolated of the radicals will have very long relaxation times due to the distance dependence of the dipolar interaction and clusters of several radicals will have very short relaxation times from cross relaxation within the clusters.^*4*^

**Figure 4.**
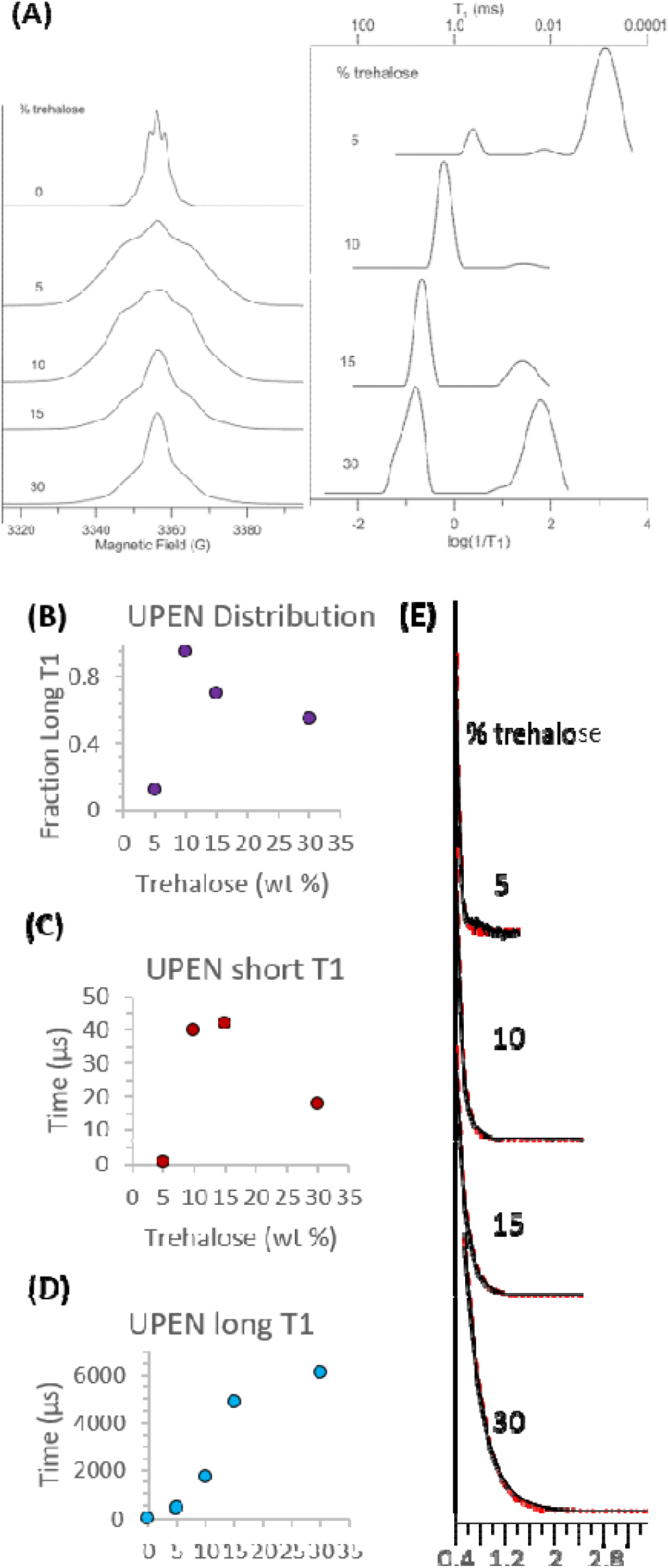
**(A)** EPR absorption spectra at 5 K for 15 mM OXo63 in aqueous samples containing 0, 5, 10, 15,or 30 % by weight trehalose. **(B)** Probability distributions of T_1_ (ms) for 15 mM solutions of OXo63 in water:trehalose mixtures calculated using UPEN from inversion recovery curves at 5K. **(C and D)** Distribution of short and long T_1_ times as function of the trehalose concentration from UPEN analysis. **(E)** Two pulse echo decays as a function of trehalose concentration. Black is the experimental data, red is the fit with single exponentials.

Changes in trehalose concentration resulted in dramatic changes in both the magnitude and distribution of relaxation times (Figure 4) of Oxo63. Without trehalose, the sample yielded an FID which could not be suppressed with attempts to decrease the magnetic field homogeneity and a short T_1_ (ca. 6-7 μs). This was also the narrowest spectrum and discrete structure was evident in the form of dipolar sidebands. A narrow linewidth and short T_1_ relaxation time are both consistent with the water crystallizing upon freezing, causing high local concentrations of OXo63 as the radical is preferentially excluded from the newly formed ice crystal. The echo-detected lineshapes of the samples containing trehalose are considerably broader, likely reflecting a more inhomogeneous environment. Inhomogeneity was evident in the relaxation as well – all of the trehalose samples exhibited wide ranges (orders of magnitude) in T_1_ relaxation times. Multiple choices of time windows therefore had to be measured to include both the short relaxation times and the long relaxation times. Values of T_m_ were short, consistent with the high Oxo63 concentrations, and echo decays could be fit well with single exponentials.

At 5% trehalose, where DNP efficiency is similar to samples without trehalose (Figure 1), the T_1_ relaxation is dominated by a fast relaxing population with a very short relaxation time (ca ~0.8 μs)^*4*^ (Figure 4a) similar to that found in the trehalose-free sample (data not shown). Two other minor populations, a distribution centered on 450 μs (10%) and another at 90 μs (~ 2%), were also detected in addition to the fast relaxing species. Increasing concentrations of trehalose caused both a population shift towards the slower relaxing species and within the slower relaxing population, a shift towards slower relaxation times. At 10% and 15% trehalose, where DNP efficiency is close to optimal (Figure 1), the long relaxing species is dominant (95% and 70%, respectively). The highest concentration, 30%, partially reverses this trend as there is a large shift in the distribution back to short relaxation times (55%). In contrast to the complex concentration dependence of the T_1_relaxation times, the linewidth decreases and T_m_ increases modestly and monotonically with the trehalose concentration. Overall, as expected, there is a rough correspondence between the T_1_ relaxation times and DNP build up kinetics. In this case, longer relaxation times are associated with faster build up kinetics, as expected for narrow linewidth radicals like Oxo63 where rapid spin diffusion exists and the solid effect is an important component in DNP transfer.

## Discussion

Solutions of trehalose have several potential advantages over the standard concentrated glycerol protocol. While the equilibrium polarization is lower in the urea test case, the build-up rate is higher and the actual polarization is similar on the 15-30 minutes time scales relevant for clinical use. Developments in coil design,^*30*^ signal acquisition,^*31*^ and post-processing,^*32*^ have significantly increased SNR since the inception of DNP MRI. The polarization process, however, is constrained by fundamental physics and will likely remain a choke point for clinical translation, especially if capital costs constrain work to a few research centers.^*2*^ The ability to rapidly polarize a sample will likely be useful in this context.

Trehalose also has the advantage of being essentially biologically inert when injected intravenously. The extent to which glycerol’s potential perturbation of metabolic activity is a concern will depend strongly on the pathway studied, but there is evidence that an acute challenge can alter a competing reaction even within the rapid timescale of the DNP experiment. Of particular concern is the rapid consumption of cytoplasmic ATP and FAD that an infusion of glycerol can cause by feeding the glycerol-3-phosphate shuttle. Marcos-Rius et al showed that a bolus of fructose several minutes before an injection of hyperpolarized ^13^C DHAP caused a significant drop in labeled G3P and PEP resonances due to the depletion of the ATP and NAD pool.^*33*^ Given that there are many metabolic reactions that are dependent on either ATP or are redox sensitive, this may be a concern as the pool of hyperpolarizable substrates continues to expand.

Several alternative methods have been proposed to improve polarization efficiency by preventing phase separation of the radical during the freezing. Lama et al modified the rate of freezing by freezing the samples in liquid-nitrogen cooled isopentane, rather than directly in liquid nitrogen, as is typically used.^*34*, *35*^ The result is a homogenous frozen solid that polarizes rapidly and efficiently regardless of the glassing properties of the original solution. The disadvantage for clinical applications is the possibility of residual traces of isopentane in the product which are not removed by the filtering process and cannot be assayed easily. Phase separation can also be prevented by impregnating the radical in a solid matrix (HYPOs and related methods),^*36*, *37*^ but the final polarization is currently less than a purely liquid formulation. Optimization of polarizing conditions will likely play a key role in the expansion of DNP MRI in the future.^*5*, *38*^

